# A novel anti-cancer therapy with nuclear export inhibitor Selinexor in combination with oncolytic myxoma virus

**DOI:** 10.1101/2022.10.31.514585

**Authors:** Masmudur M. Rahman, Fleur van Oosterom, Junior Ayuk Enow, Maksuda Hossain, Ami D. Gutierrez-Jensen, Mackenzie Cashen, Anne Everts, Kenneth Lowe, Jacquelyn Kilbourne, Juliane Daggett-Vondras, Timothy L. Karr, Grant McFadden

**Author notes:** Corresponding authors: Masmudur M. Rahman, Grant McFadden.

## Abstract

Oncolytic viruses exploited for cancer therapy are developed to selectively infect, replicate, and kill cancer cells to stop tumor growth. However, in some cancer cells, oncolytic viruses are often limited in completing their full replication cycle, making progeny virions, and/or spread in the tumor bed due to the heterogeneous cell types within the tumor bed. Here we report that nuclear export pathway regulates oncolytic myxoma virus (MYXV) infection and cytoplasmic viral replication in a subclass of human cancer cell types where virus replication is restricted. Inhibition of CRM1/XPO-1 nuclear export pathway with nuclear export inhibitors can overcome this restriction by trapping restriction factors in the nucleus and allow significantly enhanced virus replication and killing of human cancer cells. Furthermore, knockdown of CRM1/XPO-1 significantly enhanced MYXV replication in restrictive human cancer cells and reduced the formation of anti-viral granules associated with RNA helicase DHX9. Both *in vitro* and *in vivo*, we demonstrate that the approved CRM1 inhibitor drug Selinexor enhances the replication of MYXV and cell killing of diverse human cancer cells. In the xenograft tumor model in NSG mice, combination therapy with Selinexor plus MYXV significantly reduced tumor burden and enhanced the survival of animals. Additionally, we performed global scale proteomic analysis of nuclear and cytosolic proteins in human cancer cells to identify the host and viral proteins that are upregulated or downregulated by different treatments. These results for the first time indicate that Selinexor in combination with oncolytic MYXV can be used as potential new anti-cancer therapy

## Introduction

Oncolytic viruses (OVs) have emerged as novel anti-cancer immunotherapies for treating standard therapy resistant and metastatic cancers (1–3). An ideal replication-competent OV is expected to selectively infect, replicate, and generate progeny virions in the infected cancer cells, which subsequently will infect the neighboring cancer cells in the tumor bed (4, 5). This successful replication of OVs is thought to mediate antitumoral activity in multiple ways, such as the direct killing of infected cancer cells, exposing and presenting novel tumor specific neoantigens, activation of systemic antitumor and antiviral immunity, and recruitment of activated immune cells to the tumor microenvironment (TME) (6, 7). In addition to its own multi-mechanistic antitumor activity, OVs can be combined with most of the currently approved cancer therapeutics, such as chemotherapies, immune checkpoint inhibitors and cell therapies, for additional therapeutic benefits (8–11).

Myxoma virus (MYXV) has been developed as an OV against diverse malignancies (12, 13). MYXV is the prototypic member of the Leporipoxvirus genus of the Poxviridae family of viruses. Different isolates of MYXV cause disease only in European rabbits but are completely non-pathogenic to all other non-leporid species, including mice and humans. However, MYXV can productively infect many (but not all) classes of human cancer cells originating from different tissues, both *in vitro* and *in vivo*. This natural and selective tropism of MYXV for cancer cells and tissues allowed exploiting MYXV as oncolytic virotherapeutic in several preclinical cancer models for various cancer types such as pancreatic cancer, lung cancer, glioblastoma, ovarian cancer, melanoma, and hematological malignancies (12–14).

Like other poxviruses, MYXV can promiscuously bind, enter and at least initiate infection in a broad diversity of cancerous and non-cancerous cells from most vertebrate species. However, the ability of MYXV to productively replicate and produce progeny in any cell type outside the rabbit largely depends on whether the virus can successfully overcome the diverse intrinsic and innate anti-viral cellular barriers (15). These barriers are sufficiently robust to restrict MYXV replication post-entry in normal primary somatic human or mouse cells, but they tend to become compromised when cells are immortalized, transformed or cancerous. Thus, unlike rabbit cells, where MYXV can counteract every aspect of those cellular barriers, in non-rabbit normal cells and a subset of cancer cells, MYXV’s complete replication can be restricted at a different level by multiple factors. In human cancer cells, activation of these intrinsic cellular restriction factors and virus-induced signaling pathways can limit the replication and oncolytic ability of MYXV in specific cancer cell types that we refer to as either nonpermissive or semi-permissive. Several cellular pathways that are currently known to contribute to MYXV’s ability to replicate in human cancer cells include: i) endogenously activated protein kinase B (PKB)/AKT, ii) antiviral pathway activated by protein kinase R (PKR), iii) status of tumor suppressors such as p53, retinoblastoma (Rb), and ataxia-telangiectasia (ATM), iv) antiviral states induced by interferons (IFNs) or tumor necrosis factor (TNF) (16–19). In addition to these cellular barriers, we recently reported that members of the cellular DEAD-box RNA helicase superfamily have potent antiviral and/or proviral functions that regulate MYXV replication in diverse human cancer cell types (20). Among these antiviral RNA helicases, we also reported that RNA helicase A (RHA) or DHX9 exits the nucleus in response to MYXV infection to form unique antiviral granules in the cytoplasm of infected human cancer cells. These antiviral granules are formed during the late replication phase of MYXV, which reduces MYXV late protein synthesis and limits MYXV ability to replicate and generate progeny virions (21). Furthermore, DHX9 knockdown can significantly enhance the replication of MYXV in both semi-permissive and nonpermissive human cancer cell lines.

Here, we report that inhibition of the nuclear export pathway in diverse human cancer cell types where MYXV replication is restricted, significantly enhances virus replication and progeny virus formation by reducing the appearance of cytoplasmic antiviral granules. FDA-approved nuclear export inhibitor drug Selinexor also significantly enhanced MYXV replication in diverse human cancer cells. A combination of Selinexor and MYXV treatment significantly reduced cancer cell proliferation and enhanced cell killing. Furthermore, using 3D spheroid culture of human cancer cells, we show that Selinexor enhances MYXV replication and penetrative spread in spheroid cultures of cancer cells. We next tested human cancer cell derived xenograft (CDX) models in NSG mice to test the *in vivo* effect of Selinexor on oncolytic MYXV replication. Like *in vitro*cultures, Selinexor enhanced MYXV gene expression and replication in Colo205 and HT29 cells derived CDX models in NSG mice. In addition, using PANC-1 cells derived CDX models, we show that Selinexor plus MYXV treatment significantly reduced tumor burden compared to the control or MYXV treatments. Furthermore, Selinexor plus MYXV treatment significantly enhanced survival of mice. These results suggest that Selinexor and oncolytic MYXV can be developed as a novel combination therapy for cancer therapy.

## Results

### Nuclear export inhibitors enhance MYXV replication in restricted human cancer cell lines by reducing the formation of DHX9 anti-viral granules

We reported that MYXV infection of human cancer cells results in the formation of cytosolic anti-viral granules composed of RNA helicase DHX9 and reduces MYXV replication (21). Knockdown of DHX9 significantly enhanced MYXV replication and progeny virus production in cancer cells where virus replication is restricted. In uninfected cells, DHX9 is mainly localized in the nucleus; however, in MYXV infected cells, DHX9 was detected in the cytoplasm associated with anti-viral granules. Nuclear export and import pathways play a major role in the localization and function of many cellular proteins like RNA helicases (22, 23). Here we tested whether nuclear export inhibitors that target CRM1/XPO1 mediated nuclear export pathway can block the formation of the DHX9 anti-viral granules in the cytoplasm. We initially checked the effect of Leptomycin B (LMB) on MYXV replication in restricted human cancer cells such as PANC-1 (pancreatic cancer cell line) and HT29 (colorectal cancer cell line). In these cell lines, pretreatment with a lower concentration of LMB (between 0.1-0.001 μM) that had reduced cytotoxicity significantly enhanced viral gene expression, as observed with increased early/late GFP and late TdTomato reporter proteins expression (Fig 1A and 1B). This increased viral protein expression also significantly enhanced progeny virus production as measured by virus titration assay (Fig 1C and 1D). Apart from LMB, we observed a similar effect on MYXV replication with related selected inhibitors of nuclear export (SINEs) such as Leptomycin A, Ratjadone A, Anguinomycin A (data not shown). We next checked whether this increase in viral gene expression and progeny virus formation correlated with the reduction/inhibition of the formation of DHX9 anti-viral granules. Immunofluorescence staining of cells with anti-DHX9 antibody reveals that in most LMB plus MYXV treated cells (>80%), DHX9 remained in the nucleus and blocked the formation of anti-viral granules (Fig 1E and 1F).

**Figure 1.**
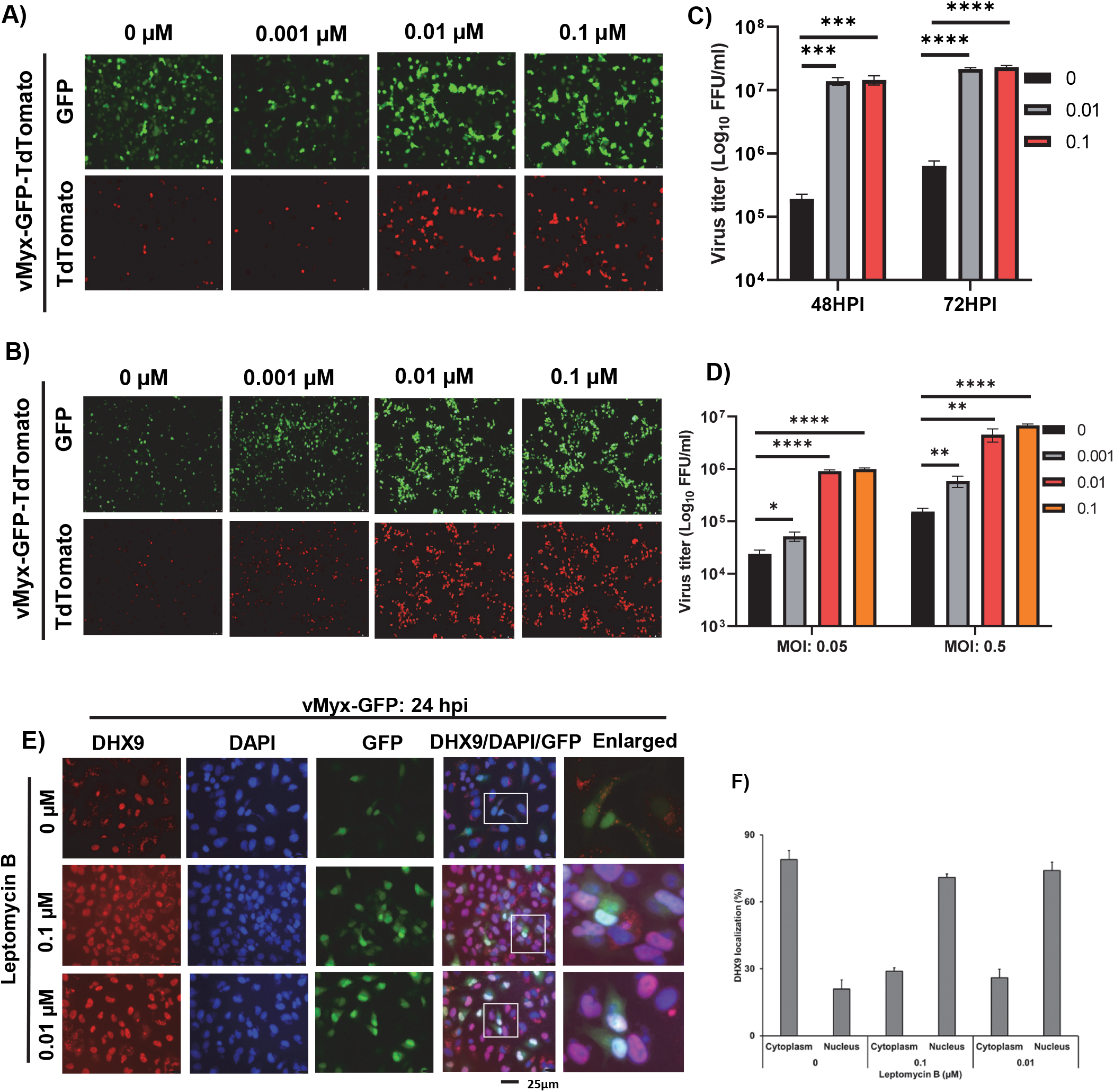
Leptomycin B treatment significantly enhanced MYXV replication in human cancer cells by reducing the formation of DHX9 antiviral granules. A) PANC-1 and B) HT29 cells were treated with different concentration of Leptomycin B for 1h, infected with different MOI of vMyx-GFP-TdTomato for 1h and replaced with fresh media containing the same doses of Leptomycin B. Fluorescence images were taken 48 hpi. C) PANC-1 and D) HT29 cells were harvested at different time points to determine progeny virus production by titration assays on permissive RK13 cells. The assays were done in triplicate. Statistically significant differences in comparison to infection without Leptomycin B are indicated. E) A549 cells were seeded on glass bottom 35mm petri dishes. Next day cells were treated with the indicated concentration of Leptomycin B for 1h and infected with vMyx-GFP (MOI = 1.0). After 24 hpi the cells were fixed and stained with antibodies against DHX9. Nuclei were stained with DAPI and imaged by fluorescence microscope. D) Number of cells showing strong nuclear or cytoplasmic staining of DHX9 after 24 hpi. A minimum of 100 cells were used for analysis from fluorescence images taken in E.

### Selinexor enhances MYXV replication in restricted human cancer cell lines

Nuclear export inhibitor Selinexor has been developed as a less toxic SINE compound to inhibit the XPO1/CRM1 mediated nuclear export pathway (24). Selinexor is approved for use in patients against hematological malignancies (25, 26). To assess whether selinexor enhances MYXV replication like LMB, multiple MYXV restricted human cancer cell lines such as PANC-1 (Fig 2A), Colo205 (Fig 2B), MDA-MB435 (Fig 2C) and HT29 (not shown) were first treated with different concentration of selinexor and infected with vMyx-GFP-TdTomato to monitor viral gene expression and replication. We also monitored and measured cell viability (described in the next section). Between 10 and 0.01 μM of Selinexor pre-treatment, we observed increased viral early/late GFP and late TdTomato reporter proteins expression in all the tested cell lines (Fig 2A, 2B and 2C). Selinexor also enhanced virus spread and foci formation in these restricted human cancer cell lines when infected at lower MOI (data not shown). However, with a concentration of 10 μM or more, Selinexor alone caused enhanced cell death in all the cancer cell lines we tested. We collected these infected cells at different time points to further assess progeny virus formation and performed virus titration using permissive rabbit RK13 cells. In all the cell lines tested, we observed a significant increase (between 1-2 logs) in virus production than infection with MYXV alone (Fig 2D, 2E and 2F). These results show that Selinexor enhances MYXV gene expression, replication and progeny virus formation in all the tested cell lines where MYXV replication is restricted.

**Figure 2.**
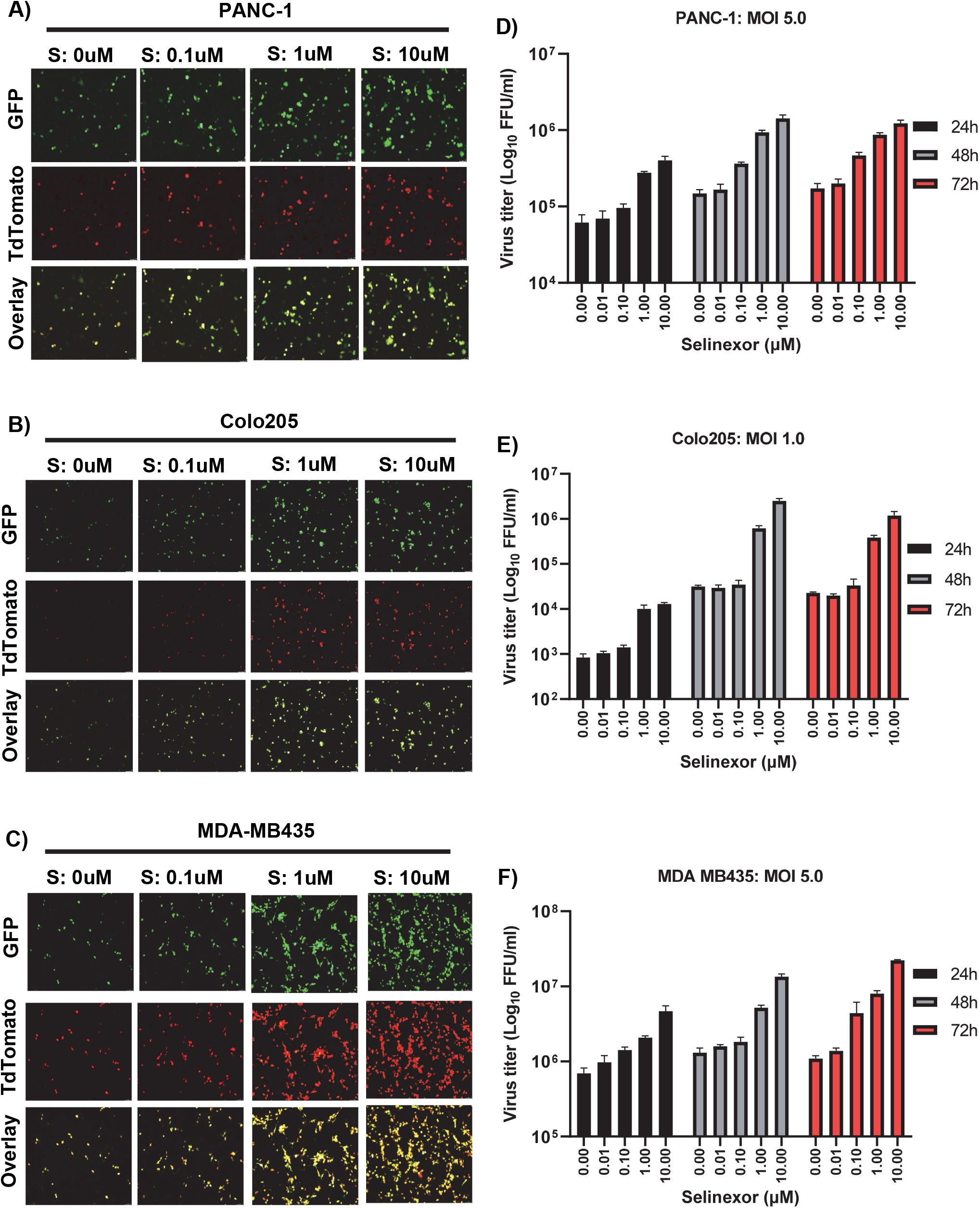
Selinexor treatment significantly enhanced MYXV gene expression and replication in diverse human cancer cell lines. A) PANC-1, B) Colo205 and C) MDA-MB435 cells were treated with different concentration of Selinexor for 1h, infected with different MOI of vMyx-GFP-TdTomato for 1h and replaced with fresh media containing the same doses of Selinexor. Fluorescence images were taken 48 hpi. D) PANC-1, E) Colo205 and F) MDA-MB435 cells were harvested at different times post infection to determine progeny virus production by titration assays on permissive RK13 cells. The assays were done in triplicate. Statistically significant differences in comparison to infection without Selinexor are indicated.

### CRM1/XPO1 knockdown enhances MYXV replication in restricted human cancer cells and reduces the formation of DHX9-containing cytoplasmic anti-viral granules

Since inhibition of the CRM1/XPO1 mediated nuclear export pathway using Selinexor or other SINEs reduced the formation of DHX9-containing anti-viral granules and subsequently enhanced MYXV replication, we further extended this observation by direct knockdown of CRM1 using siRNA. After transfection of CRM1 siRNA or control not targeting siRNA (NT-siRNA) in PANC-1 cells, infected the cells with vMyx-GFP at an MOI of 0.5 or 5.0. The level of CRM1 protein knockdown using siRNA was confirmed by Western blot analysis (Fig 3E). After CRM1 knockdown in PANC-1 cells, infection with an MOI of 0.5 allowed the formation of distinct larger foci compared to cells infected with the virus alone or NT siRNA (Fig 3A, left top and bottom panels). Furthermore, infection of PANC-1 cells at an MOI of 5.0 resulted in significantly higher GFP expression in CRM1 knockdown cells as observed under the microscope (Fig 3A, right top and bottom panels). To measure progeny virus production, we performed virus titration at 48 and 72 hpi. With both MOI of 0.5 or 5.0 we observed more than 2 log increase in progeny virus titer (Fig 3B and 3C). We then checked whether CRM1 knockdown reduced the formation of DHX9-containing anti-viral granules in the cytoplasm after MYXV infection. Immunofluorescence microscopy results demonstrated that following CRM1 knockdown, DHX9 remains in the nucleus when the cells are infected at either low or high MOI and prevent the formation of DHX9-containing cytoplasmic anti-viral granules (Fig 3D). The retention of DHX9 in the nucleus also increased the number of GFP-positive virus-infected cells in the CRM1 siRNA-mediated knockdown cells, reflecting enhanced virus infection (Fig 3D, GFP panels). These results confirm that CRM1/XPO1 nuclear export pathway is mainly responsible for the transport of proteins that form anti-viral granules and restrict MYXV replication in human cancer cells.

**Figure 3.**
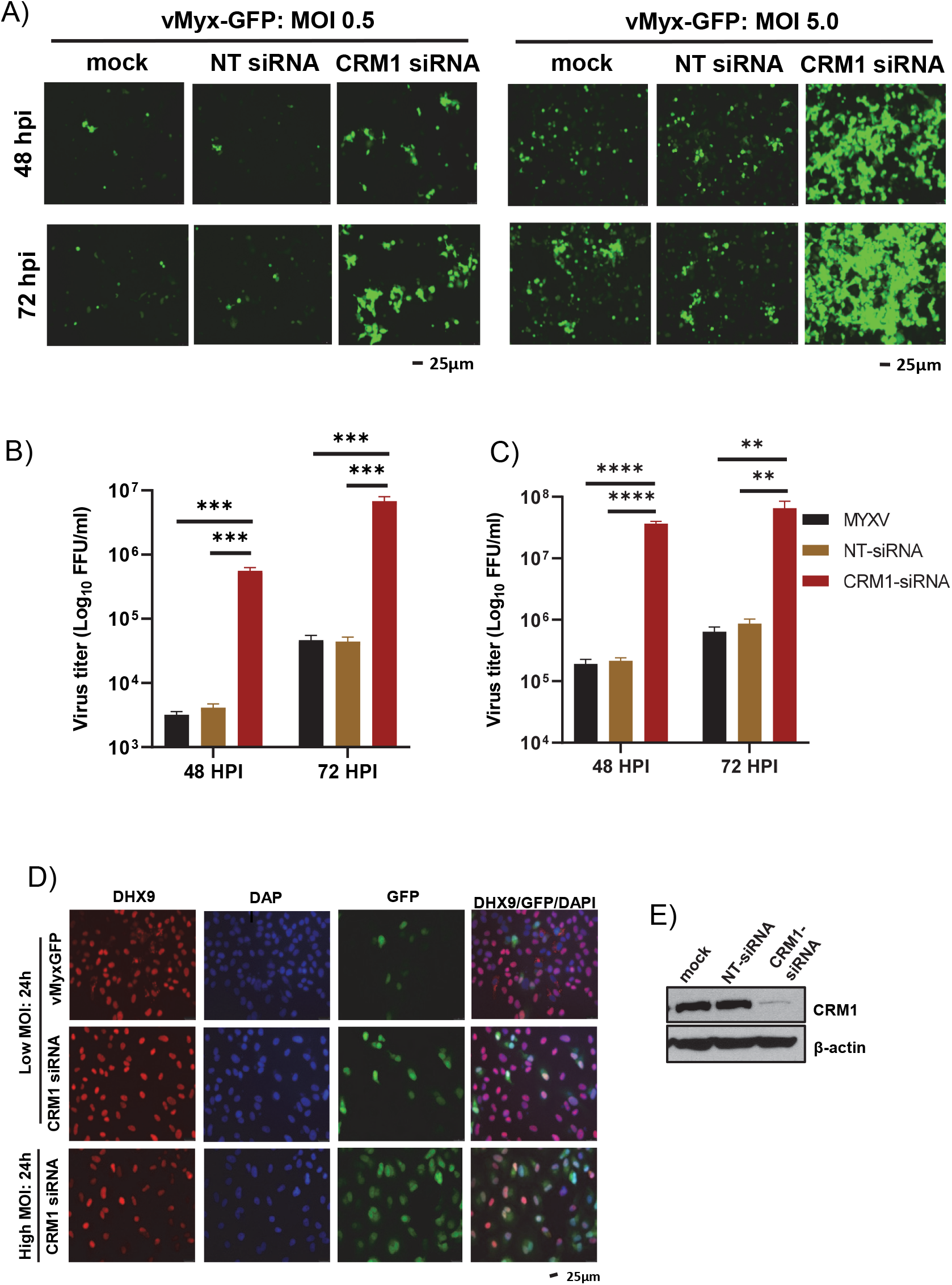
CRM1 knock down significantly enhanced MYXV replication in human cancer cells. PANC-1 cells were transiently transfected with CRM1 siRNA or control non-targeting siRNA (NT siRNA). After 48h, the cells were infected with vMyx-GFP at MOI of 0.5 or 5.0 for 1h and replaced with fresh media. A) Images showing expression of GFP after 48 and 72 hpi. B-C) The cells were harvested at 48 and 72 hpi to determine progeny virus formation by titration assay on permissive RK13 cells. Virus titers after infection with an MOI of B) 0.5 or C) 5.0. The virus titers were determined in triplicate. Statistically significant differences in comparison to infection with virus alone are indicated. D) A549 cells were seeded on glass bottom 35mm petri dishes. Next day cells were transfected with CRM1 siRNA and infected with vMyx-GFP (MOI = 0.1 or 3.0). After 24 hpi the cells were fixed and stained with antibodies against DHX9. Nuclei were stained with DAPI and imaged by fluorescence microscope. E) Western blot analysis of CRM1 protein level in A549 cells after transfection of siRNAs for 48h; actin as loading control.

### The combination of Selinexor and MYXV reduces cancer cell proliferation

Selinexor is known to reduce the proliferation of cancer cells (27). Virus infection, on the other hand, also stops cell proliferation (28). We tested this effect of selinexor and MYXV on human cancer cells when treated alone or in combination. To assess this, we performed two different cell proliferation assays. In the first method, we measured DNA synthesis by the incorporation of EdU (5-ethynyl-2’-deoxyuridine), a nucleoside analog to thymidine, into DNA during active DNA synthesis (29) (Fig 4A). Using this method in uninfected PANC-1 cells, we detected EdU incorporation in more than 50% of dividing cells (Fig 4B and 4C). When the cells were treated with Selinexor or MYXV for 24h, we observed a signification reduction in cell proliferation (20% of cells were EdU positive) than mock-treated cells. To observe the effect of a combination of selinexor and MYXV, we first treated the cells with selinexor for 1h and then infected with MYXV for 24h in the presence of selinexor. This combination treatment further significantly reduced cell proliferation to single treatment and almost completely blocked EdU incorporation (Fig 4B and 4C). Similar results were observed in other human cancer cell lines (results not shown). To further confirm these observations, we performed another cell proliferation assay using CyQUANT that measured DNA content in the cells. Using this assay, we measured cell proliferation in PANC-1 (Fig 4D and 4E) and Colo205 (Fig 4F and 4G) cells at different time points after treatment with different concentrations of selinexor and multiple MOI of MYXV. Again, in this assay, selinexor or MYXV alone significantly reduced cell proliferation after 24h compared to mock treated cells. However, the combination of selinexor and MYXV further considerably reduced cell proliferation than a single treatment at different concentrations of selinexor. These results thus suggest that the combination of selinexor and MYXV infection can almost entirely reduce cancer cell proliferation.

**Figure 4.**
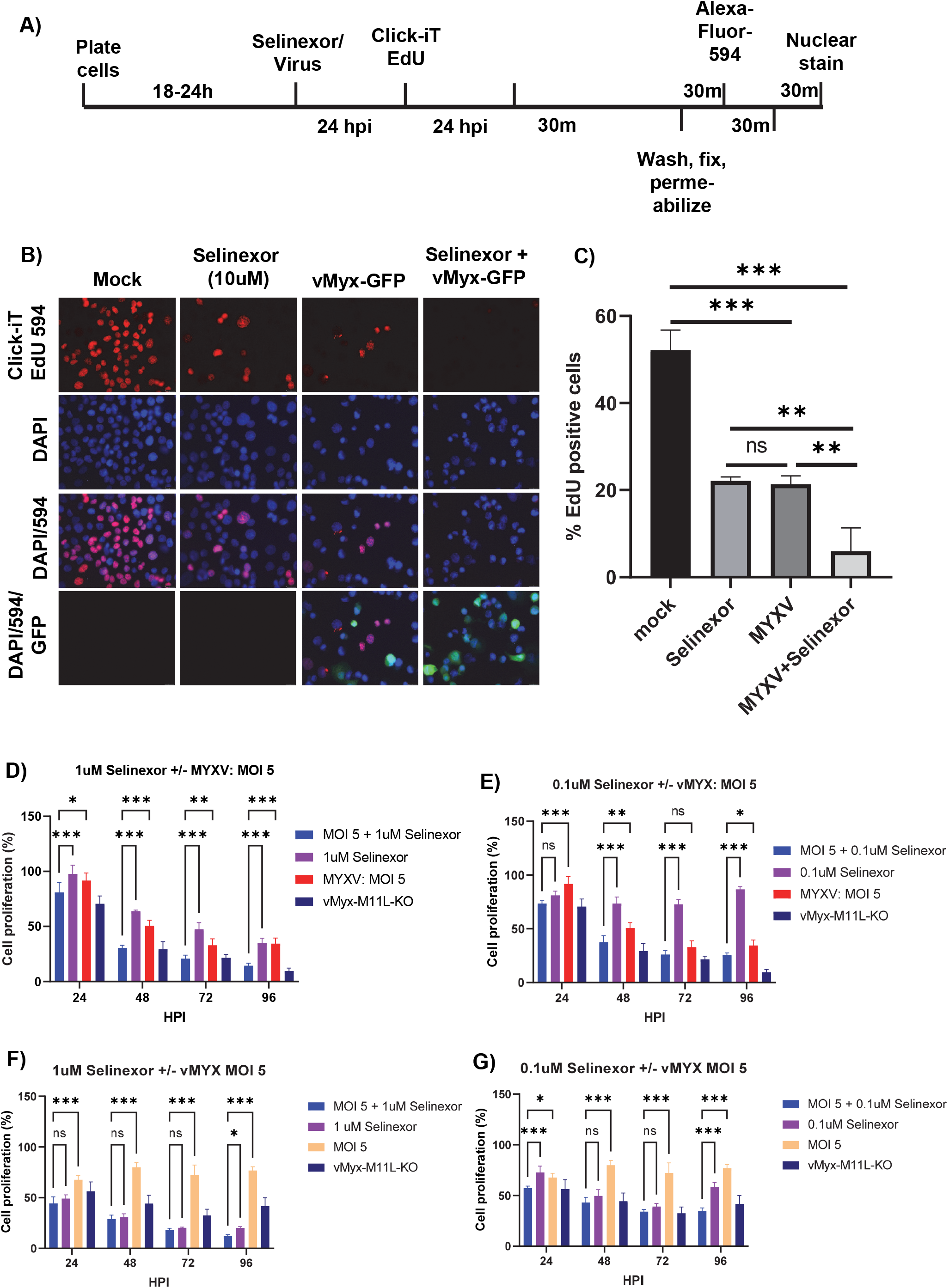
Combination of Selinexor and oncolytic MYXV significantly reduced human cancer cells proliferation. A) Schematic representation of the cell proliferation assay protocol using Click-iT EdU kit. B) PANC-1 cells were left untreated (mock) or treated with selinexor, vMyx-GFP or a combination of selinexor + vMyx-GFP and cell proliferation assay was performed using Click-iT EdU kit. The images were taken using a fluorescence microscope. C) Quantification of the number of cells showing labelling with EdU 594 (Red fluorescence). A minimum of 100 cells were used for analysis from fluorescence images taken in B. Statistically significant differences among the different treatments are indicated. PANC-1 (D-E) and Colo205 (F-G) cells in 96 well plates were left untreated (mock) or treated with selinexor, vMyx-GFP or a combination of selinexor + vMyx-GFP and cell proliferation assay was performed using CyQUANT NF (No Freeze) cell proliferation assay kit. Statistically significant differences among the different treatments are indicated.

### The combination of Selinexor and MYXV reduces cancer cell viability

To further assess whether inhibition of cell proliferation will enhance cell death, we performed cell viability assay using an MTS assay that detects metabolic activity in active cells. For this assay, PANC-1 (Fig 5A-D), Colo205 (Fig 5E-H), HT29 (data not shown) and MDA-MB435 (data not shown) cells were treated with different concentrations of Selinexor or infected with different MOI of MYXV, or treated with a combination of Selinexor plus MYXV and measured cell viability at different time points. In all the tested cell lines, when treated with 1μM or 0.5 μM selinexor, cell viability reduced to almost 50% over time; however, at lower concentrations (0.1 and 0.05 μM), selinexor had nearly no effect on cell viability. Similarly, infection only with MYXV at an MOI of 5 significantly reduced cell viability (>50%) in almost every cell line. However, MOI 1.0 and 0.5 had practically no effect on cell viability in these nonpermissive cancer cells. As described in Fig 1 and Fig 2, with these lower MOI of MYXV infection and selinexor (0.1 μM or lower) concentration, we observed enhanced viral gene expression and replication. We then tested whether combining different concentrations of selinexor treatment and MYXV infection at different MOI can further reduce cancer cell viability. Treatment with a combination of different concentrations of Selinexor and infection with an MOI of 5 of MYXV showed significantly enhanced cell death compared to selinexor or MYXV treatment alone. In particular, we observed a significant reduction in cell viability with a combination of 0.5, 0.1 and 0.05 μM selinexor and MOI 5 infection, where selinexor had almost no effect. Similarly, infection with low MOI of 1 and 0.5 and a combination of 1 μM selinexor significantly increased cell death in all the tested cell lines. These results suggest that a concentration of selinexor that does not affect cell viability can enhance MYXV replication and significantly reduce cell viability by increasing the oncolytic effect of MYXV.

**Figure 5.**
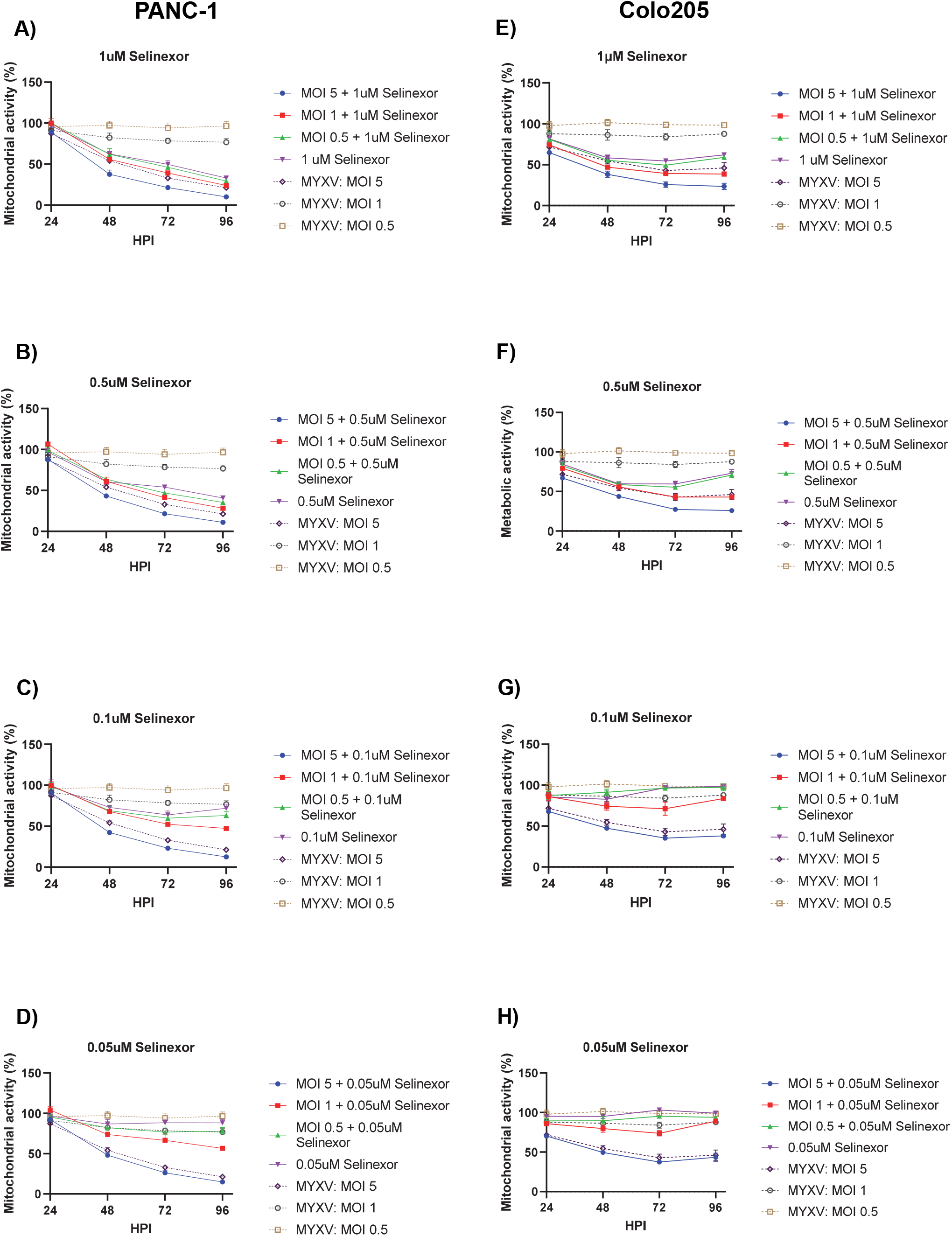
Combination of Selinexor and oncolytic MYXV significantly reduced viability of human cancer cells. PANC-1 (A-D) and Colo205 (E-H) cells in 96 well plates were left untreated (mock) or treated with different concentration of selinexor, or infected with different MOI of MYXV, or treated with a combination of selinexor and MYXV for indicated time points. Cell viability was measured using the MTS assay reagents. The assays were performed in triplicate.

### Selinexor enhances MYXV replication in 3D human cancer cell cultures

*In vitro* three-dimensional (3D) cell culture allows cell-cell to contact and forms a platform representing *in vivo* tumor mass (30). To test whether selinexor can enhance MYXV replication in the 3D culture of human cancer cells, we established a 3D cell culture using type I collagen. We used various MYXV replication-restricted human cancer cell lines such as PANC-1 (Fig 6A), HT29 (Fig 6C), MDA-MB435 (Fig 6B), and Colo205 (data not shown) to form the 3D spheroids. After forming of 3D spheroids in each well of 96 well plates, they were individually treated with different concentrations of selinexor for 1h and then infected with vMyx-GFP-TdTomato in the presence of selinexor. Images taken using fluorescence microscopy demonstrate that both GFP (early/late) and TdTomato (late) expression was enhanced in the selinexor-treated spheroids in all the tested cell lines (Fig 6A, 6C and 6E). Quantification of GFP fluorescence using a plate reader also showed a significant increase in the level of GFP expression in the selinexor treated spheroids (Fig 6B, 6D and 6F). These results confirm that Selinexor enhances MYXV gene expression and replication in 3D spheroid human cell cultures. These findings prompted us to examine the effect of selinexor on MYXV replication *in vivo* in tumors xenografted in mice.

**Figure 6.**
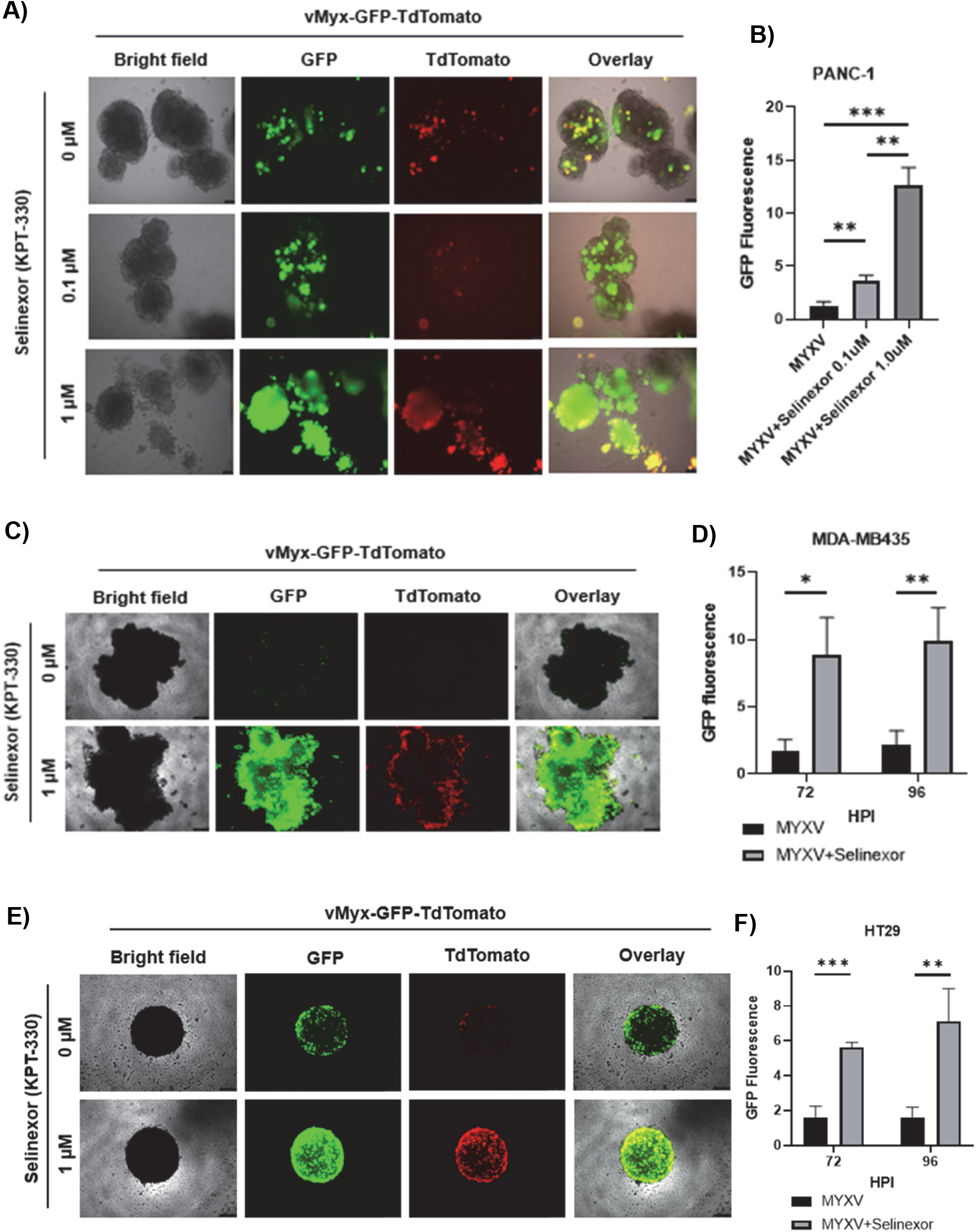
Selinexor significantly enhanced MYXV infection and replication in 3D culture of human cancer cell lines. A) PANC-1, C) MDA-MB435 and E) HT29 cells were treated with different concentration of Selinexor for 1h and then infected with vMyx-GFP-TdTomato in the presence of Selinexor. Fluorescence images were taken 72 or 96 hpi. Quantification of the level of GFP fluorescence from B) PANC-1, D) MDA-MB435 and F) HT29 cells are shown. The assays were done in triplicate. Statistically significant differences in comparison to infection without Selinexor are indicated.

### Selinexor enhances MYXV replication *in vivo* in xenografted human cancer tumors and reduces tumor burden

To test whether selinexor can enhance MYXV replication *in vivo*, we established xenograft tumor models in immunodeficient NSG mice. Human Colo205 (Fig 7) and HT29 (supplementary Fig 1) cells were injected subcutaneously on both sides of the flank to generate tumors (Fig 7A). After the tumor size reached ~100-200mm^3^, animals were randomly assigned to different treatment groups to maintain average tumor size. Animals (n=5) were than treated with either: Selinexor alone (oral), vMyx-Fluc (wild-type MYXV expressing Firefly Luc and TdTomato) alone (IT, 1×10^7^FFU on left tumor only), PBS (oral and IT) or Selinexor (oral) + vMyx-Fluc (IT, 1×10^7^FFU on left tumor only) simultaneously. After 48 h post-first treatment, mice were imaged using the IVIS system for luciferase expression. The images were analyzed, and the F-Luc signal was quantified using a software. We observed that mice that received Selinexor had significantly higher levels of luciferase signals than tumors injected with the virus alone (Fig 7B). This enhanced luciferase signal was observed in both the Colo205 (Fig 7C) and HT29 tumors (supplementary Fig 1). Subsequently, these mice received 2^nd^ and 3^rd^ dose of selinexor and IT injection of MYXV in the same tumor. IVIS imaging after the 2^nd^ treatment still showed a significantly enhanced level of luciferase signal from virus in the Colo205 tumors (Fig 7D and 7E). Additionally, we measured the tumor burden on both flanks during the course of the treatment (Fig 7F and 7G). Treatment with Selinexor alone and a combination of Selinexor plus MYXV significantly reduced tumor burden than PBS or MYXV-only treatments on both flanks in the Colo205 and HT29 xenograft models. However, when comparing the size of Selinexor and Selinexor + MYXV treated tumors, we observed no significant difference, although there was a trend that selinexor + MYXV treated tumors were smaller in size than Selinexor only treatment. These results confirmed that selinexor enhances MYXV gene expression and replication *in vivo*in animals. Additionally, Selinexor alone or combined with oncolytic MYXV can reduce tumor burden in xenograft animal models.

**Figure 7.**
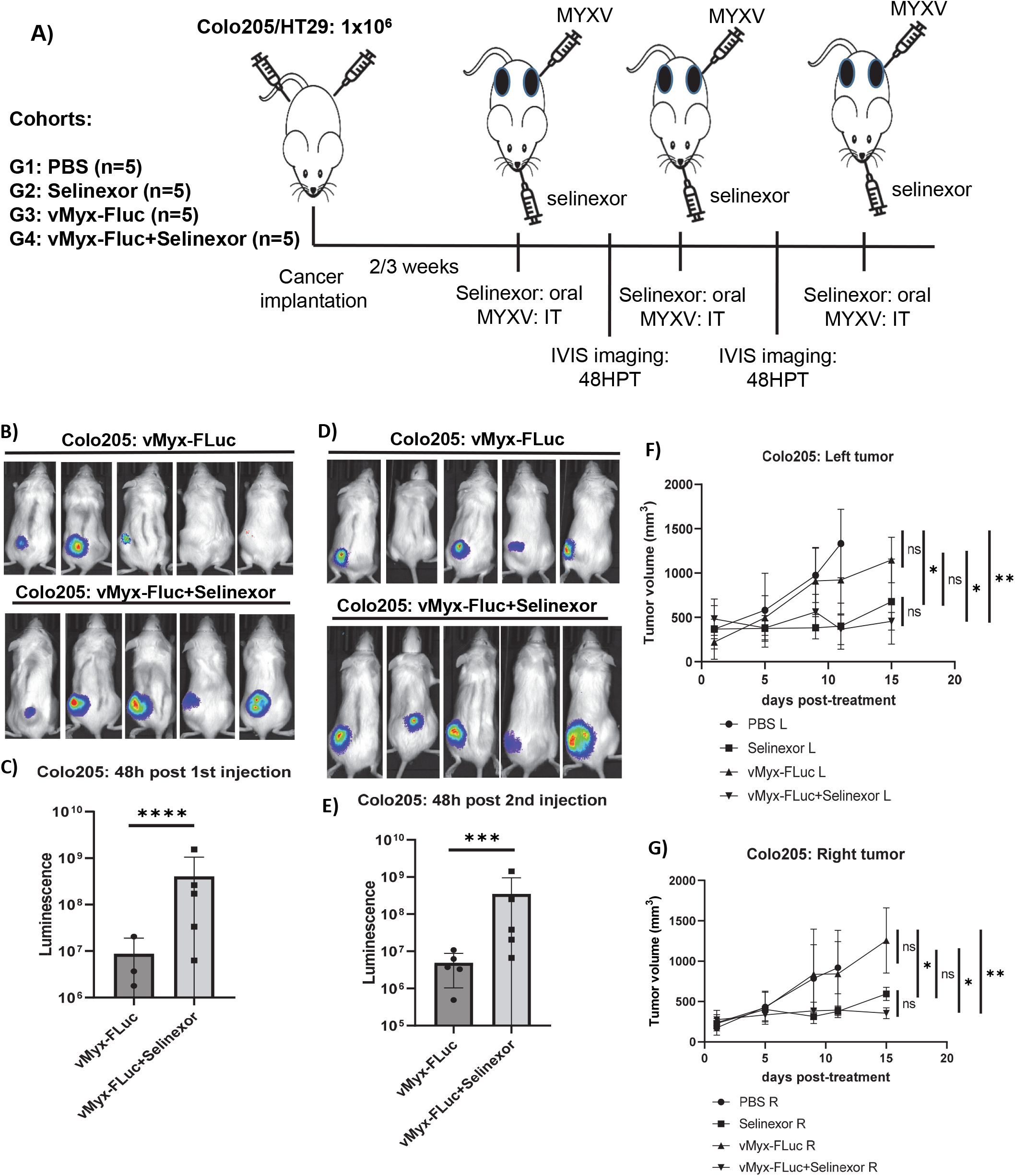
Selinexor enhances MYXV replication *in vivo* in xenograft tumors in NSG mice and reduces tumor burden. A) Diagram of experimental setup. NSG mice were inoculated with Colo205 or HT29 cells at day 0 via SQ injection on both flanks. Animals received three doses of selinexor by oral gavage and three IT injection of vMyx-FLuc on the left flank. Mice were imaged for luciferase activity 48 hours post first and second virus injection. B) Luminescence images taken using IVIS imaging system 48h after the first injection. C) Quantification of luminescence signals from images taken 48h after first injection. D) Luminescence images taken using IVIS imaging system 48h after the second injection. E) Quantification of luminescence signals from images taken 48h after second injection. F-G) Tumor volumes on the left and the right flanks were measured at different days after the start of treatments. Results are presented as mean ± SEM.

### Selinexor in combination with MYXV reduces tumor burden and extends the survival of animals in PANC-1 xenograft tumors

Based on our *in vivo* results showing that Selinexor enhances MYXV replication in the tumor bed and reduces tumor burden in Colo205 and HT29 xenograft tumors, we extended the study using PANC-1 cells xenograft tumors. Human PANC-1 cells were injected subcutaneously on both sides of the flank to generate tumors (Fig 8A). After the tumor size reached ~50-100mm^3^, animals were randomly assigned to different treatment groups such that each group maintained average tumor size. Animals (n=6) were then treated with either: Selinexor alone (oral), vMyx-Fluc alone (IT, 2×10^7^FFU on right tumor only), PBS (oral and IT) or Selinexor (oral) plus vMyx-Fluc (IT, 2×10^7^FFU on right tumor only). Animals received total 4 treatments within the first two weeks and tumor burden was measured 2-3 times every week. After the first treatment, mice were imaged using the IVIS system for luciferase expression at 24 and 72 hours post-treatment (Fig 8B and 8C). We observed that mice that received selinexor had a significantly higher level of luciferase signals than tumors injected with the virus alone (Fig 8C). After the imaging and measuring luciferase signals, we provided three additional treatments to test the therapeutic effect on the tumor burden and survival of animals. In addition, we also measured the body weight for any toxicity to the animals from the treatment (data not shown). In this PANC-1 xenograft model, we observed a significant reduction in tumor burden on both sides after treatment with selinexor alone compared to PBS or MYXV-only. Treatment with Selinexor plus MYXV significantly reduced tumor burden than PBS or MYXV-only. However, no significant difference was observed between selinexor and selinexor + MYXV treatments. In this model, again, we observed overall reduced tumor burden with the combination treatment of selinexor + MYXV than selinexor alone. When we analyzed the survival of animals from different treatment groups, animals treated with Selinexor or Selinexor +MYXV survived significantly longer than animals that were treated with either vMyx-Fluc or PBS alone (Fig 8F). We also observed that animals treated with the combination of Selinexor + MYXV survived significantly longer than animals that were treated with Selinexor alone.

**Figure 8.**
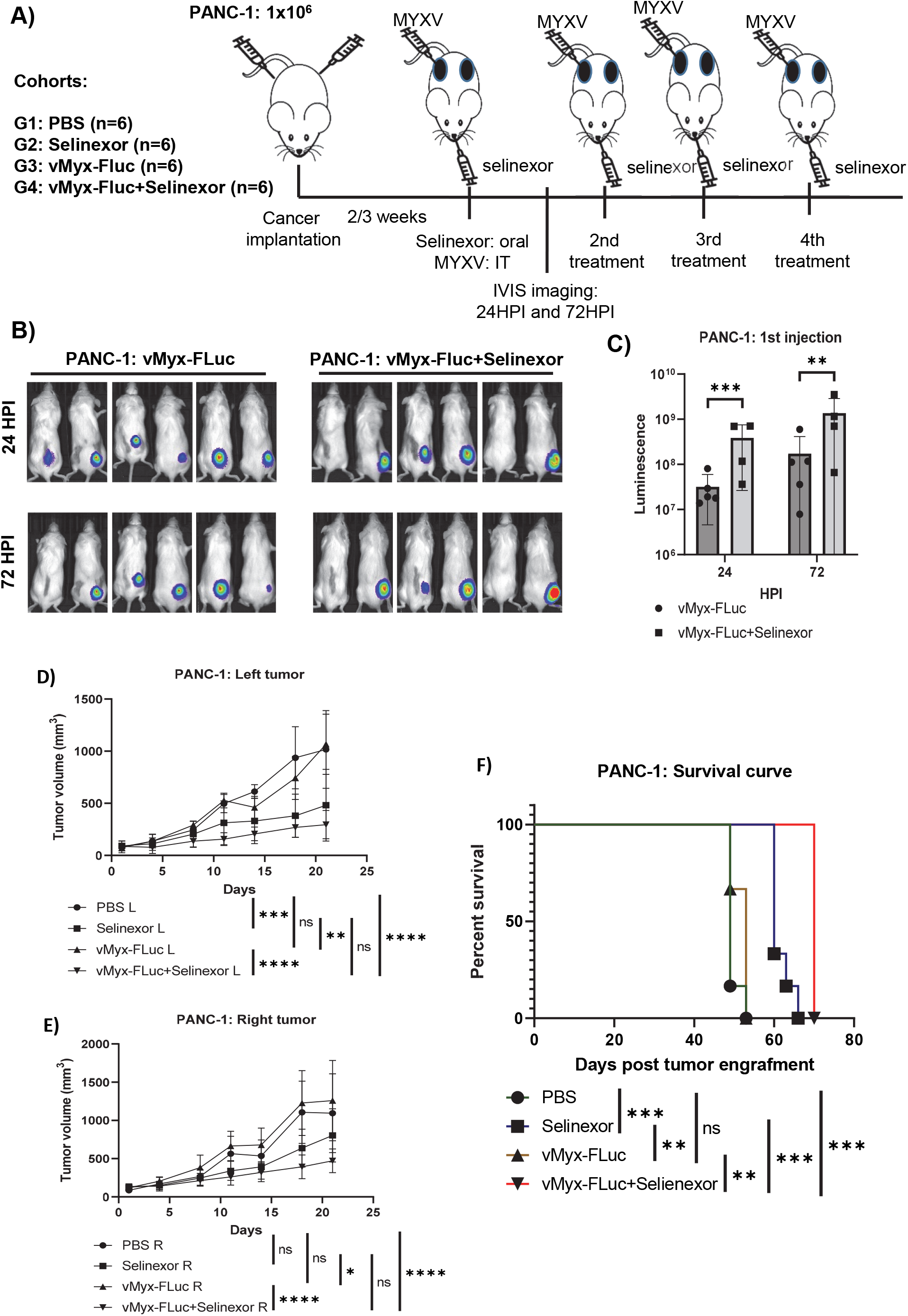
Selinexor significantly enhances MYXV replication in PANC-1 xenograft tumors in NSG mice, reduces tumor burden and prolong survival. A) Diagram of experimental setup. NSG mice were inoculated with PANC-1 cells at day 0 via SQ injection on both flanks. Animals received four doses of selinexor by oral gavage and three IT injection of vMyx-FLuc on the right flank. Mice were imaged for luciferase activity 24 and 48 hours post first virus injection. B) Luminescence images taken using IVIS imaging system 24h and 72h after the first injection. C) Quantification of luminescence signals from images taken 24h and 72h after first injection. D-E) Tumor volumes on the left and the right flanks were measured at different days after the start of treatments. Results are presented as mean ± SEM. F) Kaplan-Meier survival curves comparing animals with different treatment groups.

Additionally, we measured luciferase signals in the animals before the endpoint to confirm the presence of virus in the tumor bed after the last (fourth) injection. Mice injected with MYXV alone (10 days post last injection) showed high luciferase signals in the virus-injected tumors (Fig 9A and 9B). At this point, mice that received Selinexor + MYXV also showed a relatively higher level of luciferase signals in the injected tumors than in treatment with MYXV alone. Next, we measured luciferase signals in mice treated with selinexor + MYXV (23 days post last virus injection) when they reached the endpoint. At this point, we observed very high luciferase activity in the injected tumor and very little (one mouse) or no signal in the un-injected tumor (Fig 9C).

**Figure 9.**
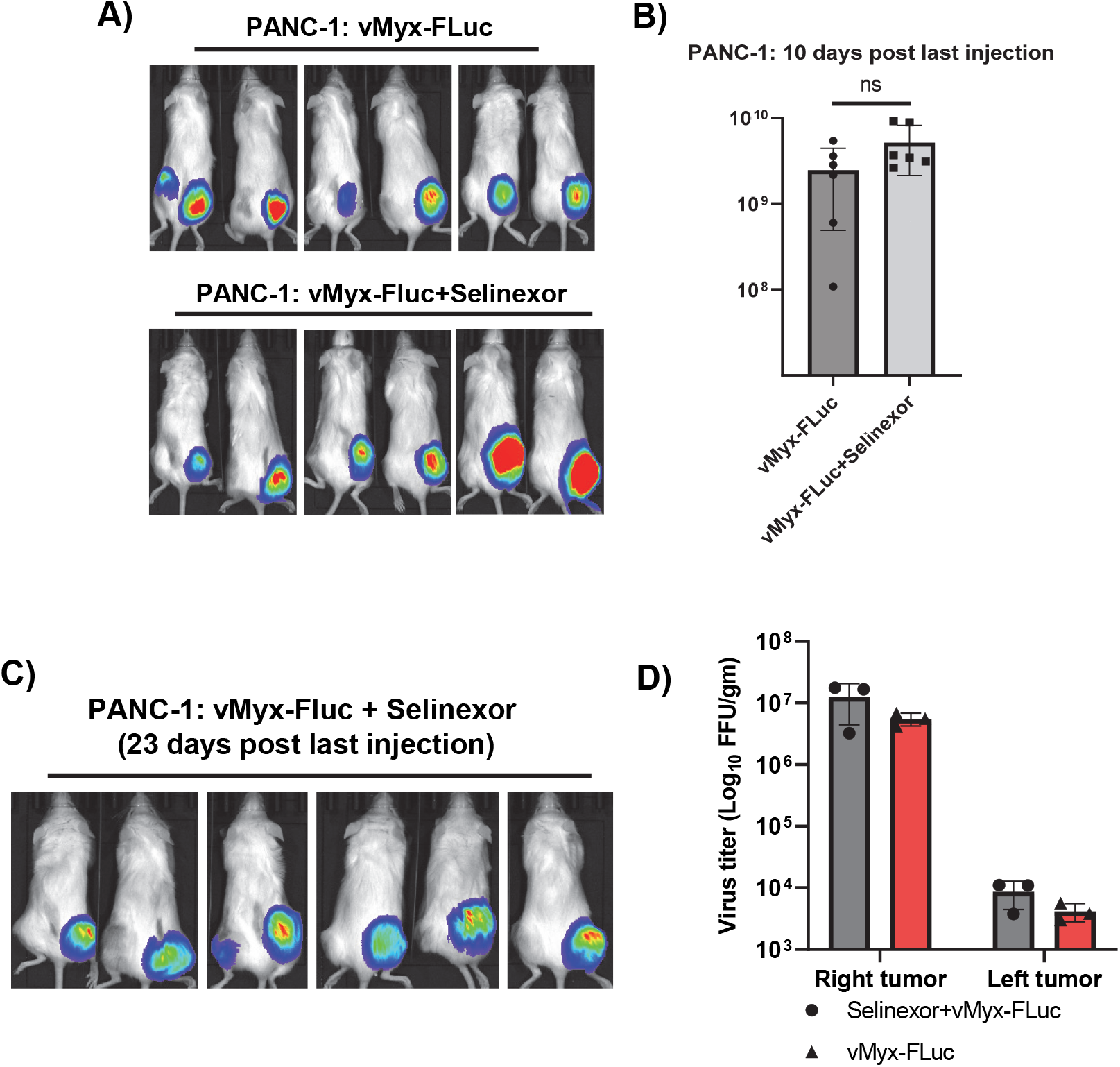
Prolonged replication of MYXV in the tumor bed of the Selinexor treated mice. NSG mice with SQ tumors of PANC-1 cells that received four doses of selinexor by oral gavage and three IT injection of vMyx-FLuc on the right flank were imaged 10 days and 23 days after the last injection of MYXV. A) Luminescence images taken using IVIS imaging system 10 days post last MYXV injection. B) Quantification of luminescence signals from images shown in figure 9A. C) Luminescence images taken using IVIS imaging system 23 days post last MYXV injection. D) Tumors collected from the mice that reached end point were processed for virus isolation and the amount of virus was titrated by foci formation assay using RK13 cells. Results shown are from three mice from each treatment groups.

Finally, we collected both the tumors from three mice of each group and performed virus titration assay (Fig 9D). To our surprise, we detected a lower level of virus from the un-injected tumor, although we didn’t detect any luciferase signals. Again, tumors that received selinexor + MYXV than MYXV alone showed a relatively higher levels of virus load than the tumors that received only MYXV.

### Both cellular and viral protein expression levels are altered in the cytoplasmic and nuclear compartments after different treatments

We performed a global proteome analysis of cytosolic and nuclear compartments to identify the cellular and viral proteins that are changed with different treatments and may contribute to enhanced virus replication, reduced cell proliferation and cell death. Human colorectal cancer cells Colo205 were used for this assay. Samples from mock and those treated with Selinexor, MYXV or a combination of Selinexor + MYXV were prepared in quadruple and processed for preparing the nuclear and cytosolic fractions. Approximately 5,000 cellular and viral proteins were identified by mass spectrometry, and their relative abundance in the nuclear and cytosolic fractions was calculated (Fig 10). We observed the most significant reduction in the abundance of proteins in nuclear and cytosolic fractions after combining Selinexor + MYXV. This is likely due to the substantial decrease in cell proliferation after the combination treatment.

**Figure 10.**
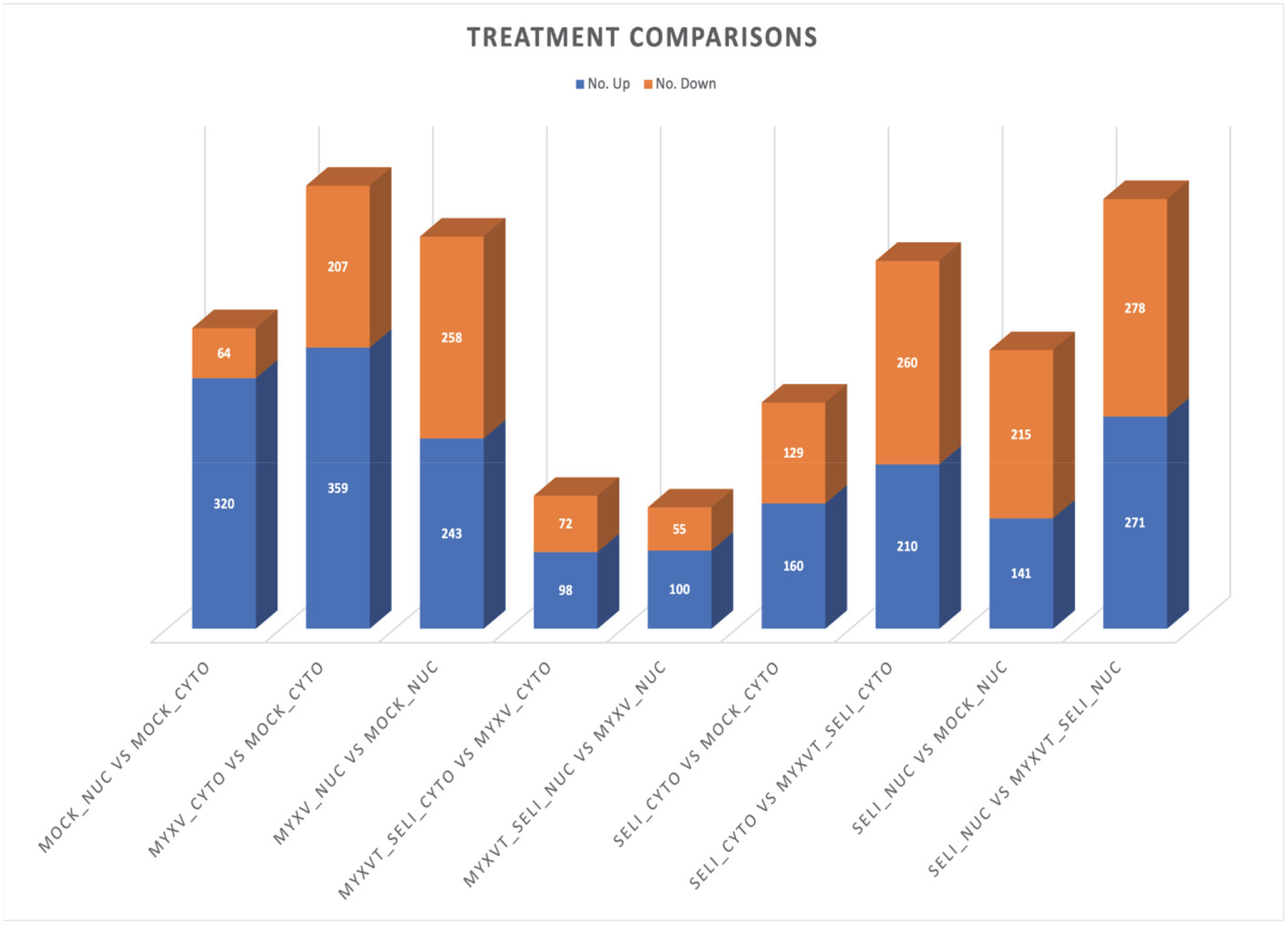
Mass spectrometry of host and viral proteins in the cytoplasmic and nuclear compartments. Human Colo205 cells were collected 48h after treatment with Selinexor, MYXV or Selinexor + MYXV and nuclear and cytosolic fractions were prepared. The assay was done with quadrupole samples from each treatment. The nuclear and cytosolic fractions were used for LC-MS analyses.

## Discussion

Cancer is the second leading cause of death, and the global number of cancer-related deaths is increasing. Therefore, novel treatment strategies are needed to improve therapeutic outcomes. Among the many new cancer treatment approaches, OVs have shown tremendous potential in preclinical animal models and clinical trials, allowing the approval of a few OVs for patients (1). However, there are still limitations with OVs that need to be addressed to get more widespread enhanced therapeutic benefits from this treatment approach. One such area of potential development is understanding how OVs and cancer cells interact. This is mainly because of the heterogeneity and the complexity of the cancer cells in the tumor bed that can alter the OV’s ability to replicate in cancer cells. Here we show for the first time that targeting nuclear export pathway can enhance the replication of oncolytic MYXV in normally restricted human cancer cells (defined as either semi-permissive or nonpermissive) and thereby enhance its oncolytic ability in preclinical animal models. Like other poxviruses, oncolytic MYXV can promiscuously bind, enter and initiate infection of most cancer cell types from different tissues and species. Still, successful productive replication that leads to progeny virus production and eventual killing of cancer cells largely depend on the viral manipulation of multiple intracellular signaling pathways (12, 31, 32). Every cancer cell has a unique spectrum of deficiencies in their cellular innate defence pathways that normally attempt to restrict virus infections, and so human cancer cells come in three general classes with respect to susceptibility to infection and killing by MYXV: fully permissive (ie produce viral progeny at levels comparable to rabbit cells), semi-permissive (ie produce at least an order of magnitude reduced levels of viral progeny) and nonpermissive (little or no viral progeny). This study refers to MYXV tropism in human cancer cells from the latter two categories.

Among the known cellular factors in cancer cells, several members of the DEAD-box RNA helicases regulate MYXV replication levels in human cancer cells (20). These RNA helicases either inhibit MYXV replication (ie antiviral) or are required for optimal virus replication (ie proviral). We recently reported that DHX9/RNA helicase A (RHA) forms unique antiviral granules in the cytoplasm that when formed inhibit MYXV replication in human cancer cells (21). DHX9 antiviral granules in the cytoplasm function by reducing viral late protein synthesis and progeny virus formation. DHX9 knockdown in restricted human cancer cells significantly enhanced MYXV gene expression and progeny virus production, cell-to-cell spread and foci formation. Apart from MYXV, DHX9 is known to also have either proviral or antiviral roles against diverse RNA and DNA viruses (33, 34). However, the diverse functions of DHX9 depend on the cell types and the localization of the protein in the infected cells.

Like many other nuclear RNA helicases, DHX9 shuttles between the nuclear and cytosolic compartments to perform cellular functions (35, 36). For example, DHX9 is imported by classical importin-alpha/beta dependent pathway (37). However, during RNA virus replication, DHX9 is also detected in the cytoplasm of the infected cells (38, 39). Based on the observations that DHX9 shuttles between nuclear and cytosolic compartments, we tested nuclear export inhibitors which target XPO1/exportin1/CRM1 to block the nuclear export of proteins in MYXV-infected human cancer cells. To our surprise, unlike RNA viruses, blocking the nuclear export pathway using the XPO1 inhibitor Leptomycin B (LMB) in human cancer cells significantly increased MYXV replication, similarly to what we observed with the knockdown of DHX9.

Additionally, LMB treatment significantly reduced the formation of DHX9 antiviral granules in the cytoplasm of MYXV-infected cells. To further confirm this enhanced virus replication XPO1/CRM1 specific, we specifically knocked down the expression of CRM1 using siRNA. Like LMB treatment, CRM1 knockdown also significantly enhanced MYXV replication in normally restrictive human cancer cells and reduced the formation of DHX9 antiviral granules in the cytoplasm. These results suggest that cellular restriction proteins exported using CRM1 have inhibitory effects on the cytoplasmic replication of MYXV. This is the first report that blocking the CRM1-mediated nuclear export pathway can enhance the replication of any virus and is opposite to what has been reported for many RNA viruses, such as HIV-1, influenza, respiratory syncytial virus (RSV), dengue virus, rabbies virus and human cytomegalovirus (HCMV), which all depend on CRM1 nuclear export pathway for replication (40).

Since LMB is relatively toxic to mammalian cells and unsuitable for *in vivo* studies in preclinical animal models, synthesized derivatives were developed and tested as potential anticancer drugs with minimal toxicity (24, 41). One such LMB derivative, Selinexor (KPT330), is approved by the FDA and is suitable for *in vivo* studies (25). Like LMB, Selinexor also significantly enhanced MYXV replication in all the human cancer cell lines tested where replication of MYXV is normally restricted. Apart from enhancing virus production, the combination of Selinexor with MYXV significantly reduced cell proliferation and enhanced cancer cell killing. More importantly, our results show that Selinexor, which has minimal toxicity to the cells, can dramatically increase virus replication and cytotoxicity against cancer cells. Thus, our results for the first time demonstrate that Selinexor can enhance the oncolytic activity of MYXV. Next, we tested whether Selinexor can enhance MYXV infection and replication in 3D organoid-like culture of human cancer cells where virus replication is restricted to the outer shell of cell spheroids. We established 3D cultures of multiple different MYXV-restricted human cancer cell lines. When treated with Selinexor and infected with MYXV, we observed a significant increase in viral early and late gene expression compared to MYXV infection alone and greater penetration into the spheroid interior. These positive results from the 3D organoid-like culture motivated us to test Selinexor and MYXV *in vivo* in animal models.

In 2019 FDA approved Selinexor for hematological malignancies such as Multiple myeloma and lymphoma (25). However, Selinexor has also shown promising results against solid tumors in preclinical animal models and clinical trials (42–44). Selinexor is delivered orally, and thus it has the potential to be combined with OV delivered either intratumorally or systemically. To test whether Selinexor will enhance MYXV replication and oncolytic activity *in vivo*, we established a xenograft model using human cancer cells implanted subcutaneously in NSG mice. Our *in vivo* studies with three different MYXV-restricted human cancer cell lines, Colo205, HT29 and PANC-1, clearly demonstrated that Selinexor significantly enhanced the replication of MYXV, as observed by measuring virus-derived luciferase signal *in situ*. To assess any therapeutic effect of MYXV, Selinexor or Selinexor + MYXV, we injected the virus intratumorally into one of two flanked tumors and delivered Selinexor systemically by oral gavage multiple times. We measured the tumor burden during the treatment; however, since there were tumors on both sides of the flank, mice had to be sacrificed when either one of the tumors reached the endpoint criteria. Selinexor alone significantly reduced tumor burden bilaterally in all the tested xenograft models compared to the PBS control or MYXV-only treatment. More importantly, treatment with Selinexor + MYXV further reduced the tumor burden in both the virus-injected and non-injected tumors compared to Selinexor alone treatment. From these studies in NSG mice, which are defective for any virus-induced acquired immunity against tumors, it was surprising that tumors that were not intratumorally injected with MYXV also showed more reduced tumor burden than those treated with Selinexor alone. To test whether MYXV was present in the un-injected tumors, we collected the tumors from the PANC-1 xenograft mice, and virus titration showed the presence of MYXV on the un-injected tumor, but only at a very lower level. At this moment, it is difficult to conclude whether the presence of migrated MYXV, innate immune cells or a combination of both contributed to this apparent abscopal tumor reduction. Another key finding is that in NSG mice, we observed persistence of the virus in the injected tumor bed for a relatively prolonged time due to the absence of an active antiviral immune system. This also contributes to the overall reduction in tumor burden, which is reflected in the PANC-1 xenograft model when we did the end-point survival study, where Selinexor + MYXV treatment significantly enhanced the overall survival of the animals. Overall, we observed enhanced therapeutic effects in mice treated with Selinexor + MYXV compared to treatment with Selinexor or MYXV only.

Finally, we performed proteomic analyses of human colorectal cancer cells Colo205 after treatment with Selinexor, MYXV and a combination of Selinexor and MYXV to determine the global expression level changes in the cellular and viral proteins in the nuclear and cytosolic compartments. Comparing the different treatments and relative abundance of proteins in the two cellular compartments, we identified both cellular and viral proteins that are upregulated or downregulated by different treatments. At this point, we have been unable to deduce any single pathway responsible for the enhanced anti-cancer activities of MYXV plus Selinexor but it will be of great interest to further evaluate the function of some of specific cellular and viral proteins in the context of virus replication, cell proliferation and cancer cell death.

---

**Supplementary Figure 1.**
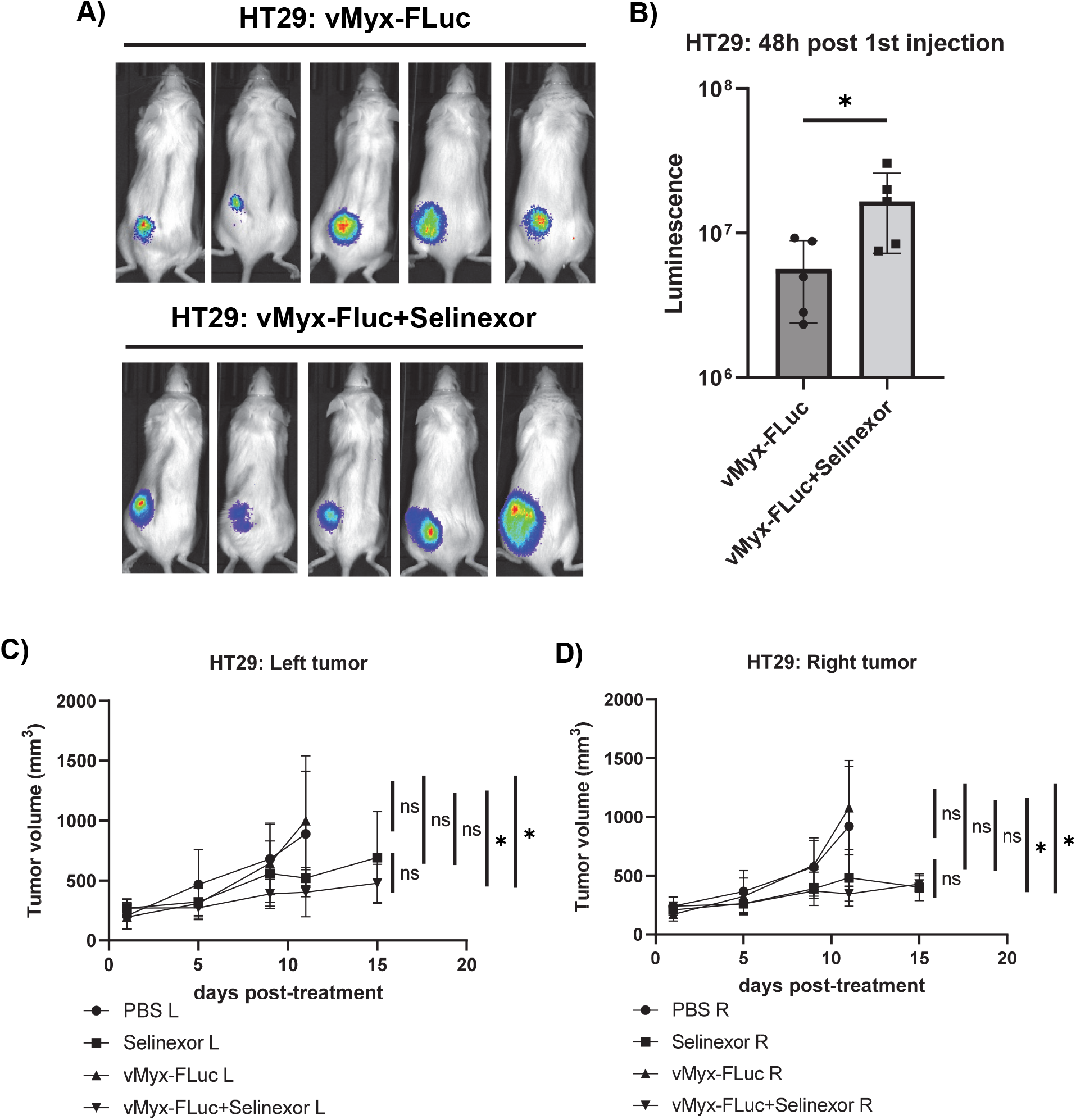
Selinexor enhances MYXV replication *in vivo* in HT29 xenograft tumors in NSG mice and reduces tumor burden. NSG mice were inoculated with HT29 cells at day 0 via SQ injection on both flanks. Animals received three doses of selinexor by oral gavage and three IT injection of vMyx-FLuc on the left flank. Mice were imaged for luciferase activity 48 hours post first virus injection. A) Luminescence images taken using IVIS imaging system 48h after the first injection. B) Quantification of luminescence signals from images taken 48h after first injection. CD) Tumor volumes on the left and the right flanks were measured at different days after the start of treatments. Results are presented as mean ± SEM.

## Materials and Methods

### Cells and viruses

Rabbit cell line RK13 (ATCC# CCL-37), non-human primate Vero cells (ATCC CCL-81), human cell lines A549 (ATCC# CCL-185), PANC-1 (ATCC# CRL-1469), and MDA-MB435 (ATCC#) were cultured in Dulbecco minimum essential medium (DMEM; source) supplemented with 10% fetal bovine serum (source), 2 mM glutamine (Invitrogen) and 100 μg of penicillin-streptomycin (pen/strep; Invitrogen)/ml. Human colorectal cancer cell line HT29 (ATCC HTB-38) and Colo205 (ATCC CCL-222) were cultured in McCoy 5 media (source) and RPMI1640 media respectively supplemented with 10% fetal bovine serum (source), 2 mM glutamine (Invitrogen) and 100 μg of penicillin-streptomycin (pen/strep; Invitrogen)/ml. All cultures were maintained at 37°C in a humidified 5% incubator. Cells were regularly checked for mycoplasma contamination using the universal mycoplasma detection kit (ATCC 30-1012K).

### Reagents and Antibodies

Rabbit polyclonal antibody for DHX9, CRM1 and mouse monoclonal antibody for β-actin were purchased from Thermo Fisher Scientific. HRP-conjugated goat anti-rabbit and anti-mouse IgG antibodies were purchased from Jackson Immuno Research Laboratories. All the secondary antibodies conjugated to Alexa Fluor 488, 594, 568 and 647 were purchased from Thermo Fisher Scientific. Selinexor (KPT330) was purchased from ApexBio. Nuclear export inhibitors Leptomycin A and Leptomycin B, Ratjadone A, and Anguinomycin A were purchased from Santa Cruz Biotechnology.

### Viruses and viral replication assay

Wild-type myxoma viruses constructs vMyx-GFP (WT-MYXV that expresses GFP under a poxvirus Synthetic early/late promoter (sE/L), vMyx-GFP-TdTomato (WT-MYXV that expresses GFP under a poxvirus sE/L promoter and TdTomato under a poxvirus p11 late promoter), vMyx-FLuc (WT-MYXV that expresses firefly luciferase under a poxvirus sE/L promoter and TdTomato under a poxvirus p11 late promoter) and myxoma virus lacking M11L gene (vMyx-M11L-KO) were used (21, 45). All myxoma viruses were grown in Vero cells. The virus stocks used were prepared using sucrose gradient purification as described before (46).

Viral titers in different human cancer cell lines were determined by a viral replication assay. Cells were seeded in a 24 well plate (2×10^5^ cell/well). Next day, cells were treated with different concentrations of leptomycin B (LMB) or Selinexor diluted in DMEM for 1h. Subsequently, MYXV was added to the cells and incubated for 1h at 37°C. After 1h, unbound virus was washed away using DMEM and DMEM with LMB or Selinexor was added to the cells again. Cells were harvested in DMEM without LMB or Selinexor at the different indicated timepoints. After harvesting the cells, they were stored in the −80°C freezer until processed. Samples were subjected to three freeze/thaw cycles and 1-minute sonification to lyse the cells and release viral particles. Afterwards, different dilutions were made in DMEM and plated on rabbit RK13 and foci were counted after 48h using a fluorescent microscope. All assays and dilutions were performed in triplicate.

### Spheroid generation and virus infection

Different cancer cell lines were grown and maintained as described in the cell culture section and used for spheroid generation within 2-5 passages. 96 well plates were prepared with Rat tail collagen I (Gibco) to form the surface for spheroid culture. On the day of cell seeding for spheroid generation, the cells were dissociated with TrypLE reagent (Gibco), neutralized the TrypLE reagent with fresh complete media, spin down the cells and pellet resuspended in fresh complete media. After making single cell suspension, cells were counted using Countess II automated cell counter and 1000 cells in 100ul volume was plated on the surface of the collagen matrix. The cells were observed daily for the formation of spheroids. After 5-7 days, when the spheroids reached the desired size they were treated with selinexor and then infected with vMyx-GFP-Tdtomato virus

### Immunofluorescence

Cells (5×10^5^-1×10^6^/dish) were seeded on glass bottom 35mm petri dishes overnight. Depending on the experiments, next day cells were either transfected with siRNA for 48h or treated with nuclear export inhibitor, MYXV or a combination of both. At different time points after treatment cells were washed with PBS 3 times, fixed with 2% paraformaldehyde in PBS for 12 min at room temperature, again washed with PBS 3 times and permeabilized in 0.1% Triton X-100 in PBS for 90 Sec at room temperature. Fixed cells were washed with PBS 3 times and then blocked with 3% BSA in PBS for 30 min at 37°C. Samples were then incubated with primary antibody (1:300 dilution) for 30 min at 37°C, washed with PBS 6 times and incubated with secondary antibody conjugated to different Alexa Fluor. After again washing with PBS 6 times, samples were mounted on glass slides with vecta shield (Vectorlabs) containing DAPI (4’,6-diamidino-2-phenylindole) to stain DNA in the nuclei and viral factory. Images were captured using a Leica fluorescence microscope.

### siRNA transfection

ON-TARGETplus SMART pool siRNAs for CRM1/XPO1 and non-targeting control (NT siRNA) were purchased from Dharmacon (Horizon Discovery). In 24 well plate cells were seeded with 40–50% confluency, left over-night for adherence and then transfected with the siRNAs (50 nM) using the Lipofectamine RNAiMAX (Invitrogen) transfection reagent. After 48 h of transfection the cells were infected with different MOI of vMyx-GFP for 1h, washed to remove unbound virus and incubated with complete media. At the indicated time points cells were either observed under fluorescence microscope to monitor and record the expression of fluorescence proteins, or harvested and processed for titration of progeny virions.

### Click-iT EdU cell proliferation assay

To visualize and measure cell proliferation, a Click-iT EdU cell proliferation (Thermo Fisher) assay was performed according to the manufacturer’s instructions. Briefly, cells (5×10^5^/dish) were seeded on glass bottom dishes and allowed to adhere by incubation overnight at 37°C. Next day cells were treated with Selinexor, MYXV or a combination of both for 24hours. Subsequently, the EdU reagent (10μM) was added and incubated for another 24 hours. To visualize EdU incorporation in dividing cells, cells were fixed with 3.7% formaldehyde in PBS and permeabilized with 0.5% Triton X-100 in PBS. Cells were then incubated with Click-iT EdU reaction cocktail with Alexa-Fluor-594 for 30 minutes at room temperature, protected from light. The cells were washed with PBS and subsequently stained with Nuclear Mask Blue stain for nuclear staining. The fluorescence images were taken using a fluorescence microscope and fluorescence signals were analyzed using ImageJ software.

### Cell proliferation assay

To measure cancer cells proliferation based on the amount of cellular DNA, a CyQuant NF Cell Proliferation Assay Kit (Invitrogen) was used according to the manufacturer’s instructions. Briefly, Panc-1, HT29, MDA MB 435 or Colo205 cells were seeded in a 96-well plate (1·10^4^ cells/well) and left to attach to the wells overnight. The next day, medium was removed and replaced with 50μL medium containing different concentrations of Selinexor (0μM-1μM). After an hour incubation with Selinexor, the virus was added in different MOIs (MOI 0.5 – MOI 5), bringing the end volume of every well up to 100μL. A 1X dye binding solution was prepared by adding 9μl of the CyQuant NF Dye reagent in 4.5ml Hank’s Balanced Salt Solution (HBSS) buffer (Invitrogen). After 24, 48, 72 and 96 hours of incubation, medium was removed from the cells and 50μL of the 1x dye solution was added to all wells. The microplate was covered to protect it from light and incubated for 30-60 minutes in a humidified 5% CO2 incubator at 37°C. Subsequently, cell proliferation was quantified by reading out fluorescence with excitation at 485nM and emission detection at 530nM in the VarioSkan Lux Microplate reader (Thermo Fisher). All conditions were performed in quadruples and normalized to mock treated cells.

### Cell viability assay

To assess viability of different human cancer cells after Selinexor treatment or MYXV infection, 10,000 cells were seeded in each well of 96-well plate. Next day, cells were either treated with different concentration of Selinexor or infected with different MOIs of MYXV or treated with different concentration of Selinexor for 1h followed by infected with different MOIs of MYXV. A minimum of four to five wells were used for each treatment conditions and untreated cells (mock) served as control. At different time points such as 24h, 48h, 72h and 96h cell viability was assessed using MTS assay.

### Animal studies

All animal experiments were performed under the Institutional Animal Care and Use Committee (IACUC) approval of Arizona State University and conformed to all regulatory standards. Male and female NSG mice were purchased from Jackson laboratory at 6-8 weeks of age. After arrival the animals were housed in the Biodesign Institute vivarium under sterile conditions. Animals were acclimatized for at least 7 days before tumor implantation or any experimental procedures. All animal handling, housing, husbandry, and experimental protocols were carried out based on the approved IACUC protocols and institutional standards. Cells (1×10^6^/mouse in 100ul of PBS) were subcutaneously injected into the flanks of NSG mice. When average tumor volumes reached 50-200 mm^3^, mice were randomized into different treatment groups. Each treatment groups had 5 or 6 animals. Tumor volume was measured 2-3 times per week: Volume =½ (length x width^2^).

When the tumor volume reached between 1.5-2.0cm3, the animals were euthanized, and tumor was collected for histology or processed for virus titration. For detection of MYXV replication in the tumor bed, luciferin was injected by IP delivery and bioluminescence images were taken (Xenogen IVIS 2000).

### Nucleus-cytoplasmic fractionation and proteomics

Human colorectal cancer cells Colo205 were collected 48h after treatment with Selinexor, MYXV infection or Selinexor + MYXV and nuclear and cytosolic fractions were prepared using NE-PER nuclear and cytoplasmic extraction reagents (Thermo Scientific). Purity of the fractions was confirmed by western blot analysis for tubulin (cytoplasmic) and histone H3 (nuclear). These fractions were used for LC-MS analyses at the Biosciences Mass Spectrometry Core Facility at Arizona State University.

### Statistical analysis

Statistical analyses were performed using GraphPad prism software. Values are represented as means + SD for at least two or three independent experiments. The paired, two-tailed Student t test was used to determine the significance between the two groups. Kaplan-Meier analysis of mouse survival was performed with GraphPad Prism software and log-rank (Mantel-Cox) test was performed to compare survival curves and to perform statistical analysis. *P* values are reported as follows: no significant (ns) *P*> 0.05, * *P* < 0.05, ** *P* < 0.01, *** *P* < 0.001, **** *P*< 0.0001.

## Author contributions

M.M.R. designed and performed experiments, analyzed data and wrote the manuscript. F.O. performed experiments and analyzed data. J.A.E., M.H., A.D.G-J., M.C., A.E., K.L., J.K., J.D-V., performed different experiments. T.L.K. analyzed mass spectrometry data. G.M. and M.M.R. received funding.

## Acknowledgements

This work was supported by National Institute of Health (NIH) grant R01 AI080607 to G.M. and M.M.R. The funders had no role in study design, data collection and interpretation or the decision to submit the work for publication.

